# Microcystin shapes the *Microcystis* phycosphere through community filtering and by influencing cross-feeding interactions

**DOI:** 10.1101/2024.09.18.613610

**Authors:** Rebecca Große, Markus Heuser, Jonna E. Teikari, Dinesh K. Ramakrishnan, Ahmed Abdelfattah, Elke Dittmann

## Abstract

The cyanobacterium *Microcystis* causes harmful algal blooms (cyanoHABs) that pose a major threat to human health and ecosystem services, particularly due to the prevalence of the potent hepatotoxin microcystin. With their pronounced EPS layer, *Microcystis* colonies also serve as a hub for heterotrophic phycosphere bacteria. Here, we tested the hypothesis that the genotypic plasticity in its ability to produce microcystin influences the composition and assembly of the *Microcystis* phycosphere microbiome. In an analysis of individual colonies of a natural *Microcystis* bloom, we observed a significantly reduced richness of the community in the presence of microcystin biosynthesis genes. A subsequent synthetic community experiment with 21 heterotrophic strains in co-cultivation with either the wild-type strain *M. aeruginosa* PCC 7806 or the microcystin-free mutant Δ*mcyB* revealed not only a tug-of-war between phototrophic and heterotrophic bacteria, but also a reciprocal dominance of two isolates of the genus *Sphingomonas* and *Flavobacterium*. In contrast, an *Agrobacterium* isolate thrived equally well in both consortia. In substrate utilization tests, *Sphingomonas* showed the strongest dependence on *Microcystis* exudates with a clear preference for the wild-type strain. Genome sequencing revealed a high potential for complementary cross-feeding, particularly for the *Agrobacterium* and *Sphingomonas* isolates but no potential for microcystin degradation. We postulate that strain-specific functional traits, such as the ability to perform photorespiration and to produce vitamin B12, play a crucial role in the cross-feeding interactions, and that microcystin is one of the determining factors in the *Microcystis* phycosphere due to its interference with inorganic carbon metabolism.

## Introduction

Massive growth events (blooms) of cyanobacteria are a seasonally recurring problem that poses a threat to human health, water quality and ecosystem services (1). The spread of these blooms is increasingly accelerated due to changing climate which results in longer stratification periods in lakes and higher temperatures across the globe (2, 3). A widespread cyanobacterial genus that tends to form harmful algal blooms (cyanoHABs) is the unicellular genus *Microcystis* which is a predominant producer of the potent hepatotoxin microcystin (MC) (4). *Microcystis sp.* form macroscopically visible colonies that are surrounded by a distinct mucus layer which serves as a nutrient-rich habitat for the heterotrophic microbiome (phycosphere) (5). *Microcystis* and its heterotrophic interactome are increasingly regarded as holobiont (5), with the heterotrophic bacteria benefiting from the dissolved organic carbon provided by the cyanobacteria and the cyanobacteria presumably taking advantage of nutrient recycling, CO_2_ production and the reduction of reactive oxygen species by their microbiome (6–9). Cultivation-dependent studies have shown that a significant number of the associated heterotrophic bacteria have a growth-promoting effect on *Microcystis* strains (10). Therefore, the environmental success of *Microcystis* blooms must be regarded as a joint effort of this association. Extensive studies of the *Microcystis* phycosphere have provided evidence of a high specificity of interactions compared to the surrounding planktonic bacterial community (6). At the same time, however, variability and succession of the microbial community is also observed, especially in different bloom stages during the season (11).

The great environmental success of *Microcystis* is increasingly associated with the genotypic and phenotypic plasticity of the genus. Although the genus has a highly similar core genome, the increasing number of sequenced *Microcystis* genomes have revealed a large pangenome with major flexible portions (12, 13). A functional trait that shows great variability among *Microcystis* strains is the inorganic carbon adaptation. Different *Microcystis* strains encode different sets of bicarbonate uptake transporters, resulting in major differences among individual strains in their adaptation to low or high availability of inorganic carbon (14, 15). Further, a large part of the flexible genome portions is dedicated to the production of different secondary metabolites, including MC but also cyanopeptolin, aeruginosin, microginin, anabaenopeptin and aeruginoguanidine (12, 16). Notably, the genotypic and chemotypic plasticity of *Microcystis* is reflected in the associated microbiome. A recent study was able to provide clear indications of a co-phylogeny through single colony sequencing and also provided evidence of phylosymbiosis (17). However, little is known about which flexible *Microcystis* traits have the greatest impact on the specificity of the interactions. There are certainly indications that the ability to produce MC has an influence on heterotrophic partners. For example, one study showed a positive correlation between the ability to produce toxins and the occurrence of α-proteobacteria of the genus *Phenylobacterium* in Lake Taihu in China. Furthermore, it was shown that isolated *Phenylobacteria* promote the dominance of the toxic wild-type strain *Microcystis aeruginosa* PCC 7806 over the non-toxic mutant Δ*mcyB* in chemostat experiments (18).

Microbial secondary metabolites are very much perceived through their role as weapons in microbial interactions or through their therapeutic potential. However, there is evidence that the ecological role of secondary metabolites goes far beyond their potential defensive role. There are studies indicating that even sub-inhibitory concentrations of secondary metabolites influence microbial growth, biofilm formation and community behavior (19, 20). In this context, the role of secondary metabolites in the structuring of microbial communities is also increasingly being discussed. In the case of MC, the defensive function is largely limited to eukaryotic organisms. A role for MC in microbe-microbe interactions is therefore currently rather discussed in connection with the specific degradation of MC and its use as a carbon source (21). However, there is extensive evidence that the ability to produce MC has an impact on functional traits that can potentially influence microbial interactions. For example, the phenotypic, proteomic and metabolomic comparison of the toxic strain *M. aeruginosa* PCC 7806 and the Δ*mcyB* mutant showed that 1) the loss of MC affects surface components such as the lectin Mvn and the filamentous glycoprotein MrpC and concomitantly the aggregation tendency of the bacteria (22, 23); 2) the loss of MC leads to a reprogramming of the carbon metabolism, especially under high light conditions, which also influences the accumulation of extracellular dissolved organic carbon in the form of glycolate (24); 3) the loss and also the addition of MC affects the subcellular localization of the CO_2_-fixing enzyme RubisCO underneath the cytoplasmic membrane, which could possibly promote direct CO_2_ assimilation from the heterotrophic bacteria (25, 26).

In the present study, we aimed to systematically investigate the influence of MC production on the composition of the *Microcystis* microbiome. To this end, we first performed single colony analyses of MC-producing and non-producing field colonies and established a synthetic community covering a broad spectrum of phycosphere bacteria employing PCR-based discrimination and 16S-rRNA amplicon sequencing strategies. We were able to show a correlation of MC production and the composition of the microbiome in both field and laboratory samples, and further provide evidence that the exudates of cyanobacteria have a discriminating effect on the growth of individual heterotrophic species. Our study supports the hypothesis that MC is one of the flexible traits that influences the specificity of interactions in *Microcystis* holobionts.

## Results

### Richness and community composition differ in MC+ and MC- Microcystis phycospheres

To analyze a possible correlation of MC and *Microcystis* phycosphere microbiomes, we isolated single colonies from a *Microcystis* bloom of Lake Zernsee near Potsdam (Figure S1). All colonies were sampled on the same day and from the same lake to minimize the influence of seasonality, abiotic environmental factors and the surrounding free-living heterotrophic community. Single colony DNA isolation and PCR analysis of the MC biosynthesis gene *mcyA* were used to discriminate individual colonies into MC+ (12; *mcyA(+)*) and MC- genotypes (17; *mcyA*(*-*)) (Figure 1A, Figure S2). Colonies yielding only weak PCR results were excluded from further analysis. Next, the microbiomes of MC+ and MC- colonies were studied by sequencing 16S-rRNA gene amplicons spanning the V3-V4 regions. 16S-rRNA sequencing and *mcyA* PCR analyses were performed with the same DNA material. Following taxonomy assignment, individual colonies could be assigned to three different *Microcystis* ASVs (> 1000 reads). Specifically, a single dominant *Microcystis* ASV was detected in 80% of the analyzed colonies (10 colonies belonged to ASV1, 12 colonies to ASV2 and 1 colony to ASV3, see Table S1), while a mixture of two different *Microcystis* ASVs was observed in 20% of colonies. A similar observation was recently made in a single colony-study of Lake Erie *Microcystis* blooms and discussed as a result of two divergent 16S-rRNA copies in single *Microcystis* genomes (27). Based on these findings, we assume that the individual colonies isolated in this study were predominantly clonal isolates. Yet, we cannot exclude the possibility that some of the colonies contained two clonal strains. We did not observe a correlation between ASV type and the presence or absence of MC. Next, we analyzed the non-*Microcystis* ASVs in single colonies. Similar to previous studies, we did not observe a ubiquitously present core microbiome. However, certain taxa showed comparatively high prevalence. Among all colonies, the predominant heterotrophic taxa detected on genus level were *Roseomonas* and *Microscillaceae Family* (76%), *Vibrio* (72%), *Pelomonas, Tabrizicola* and *Cutibacterium* (69%), and *Phenylobacterium* and *Flavobacterium* (62%) (Figure 1A and Table S2). To evaluate possible differences between MC+ and MC- microbiomes we compared co-presence networks for the two subgroups. Both networks were found to be principally similar (Figure 1A). A total of 20 nodes were assigned to genera that are shared between the two types. However, there were also some noticeable differences. Overall, the total number of non-*Microcystis* nodes was lower in the MC+ type (24 nodes) than in the MC- type (34 nodes). Considerable differences were observed for the class Alphaproteobacteria (11 nodes (MC-), 7 nodes (MC+)) followed by Gammaproteobacteria (10 nodes (MC-), 5 nodes (MC+)) and Bacteroidia (3 nodes (MC-), 5 nodes (MC+)). The Bdellovibrionia class was uniquely represented by the *Peredibacter* genus in co-presence in the MC+ type (1 node). In MC+ colonies, two *Microcystis* ASVs showed co-presence connections, while in MC- colonies only *Microcystis* ASV1 showed co-presence links (Figure 1A).

**Figure 1.**
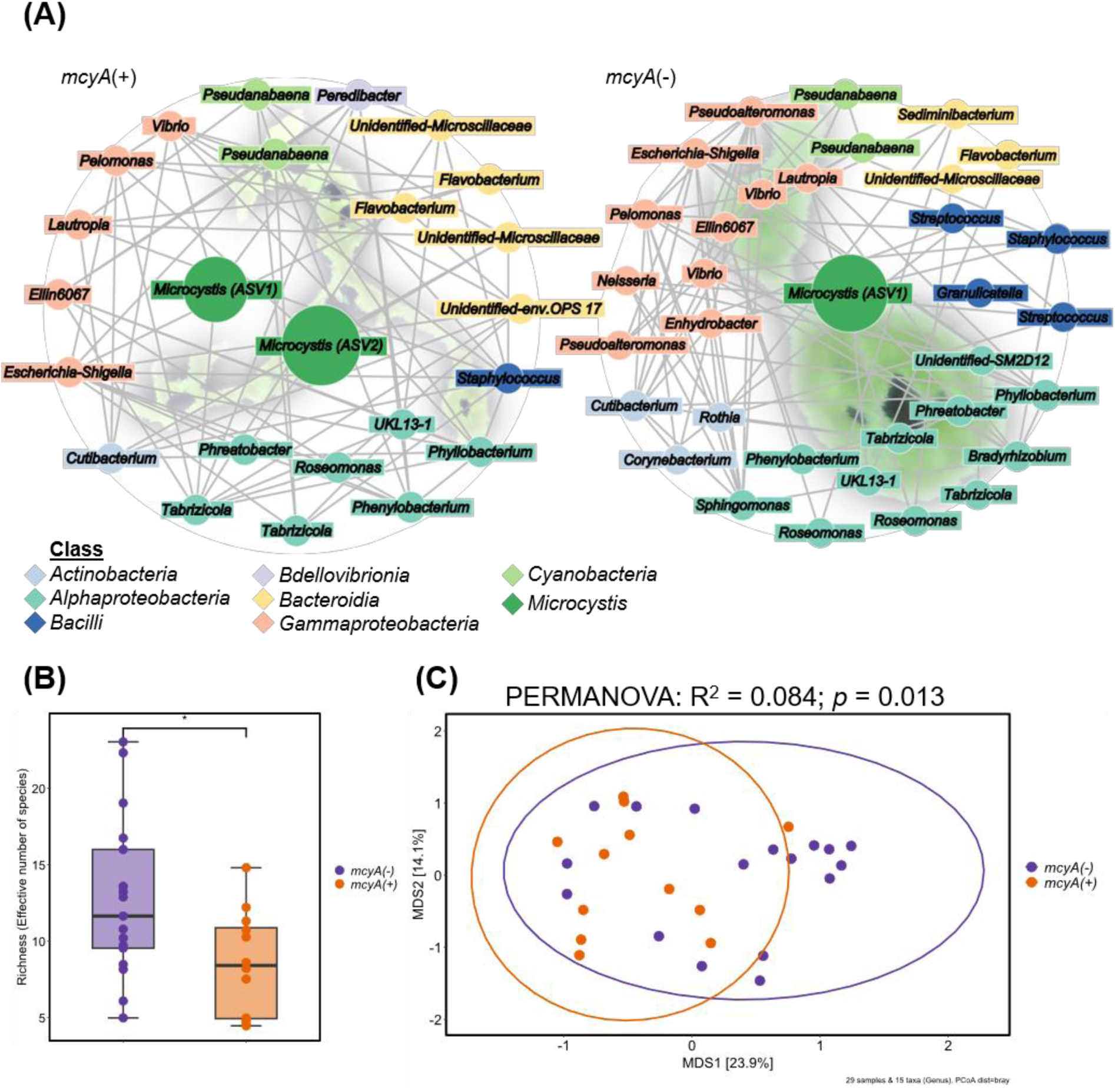
Analysis of naturally occurring microbiomes of single *Microcystis* colonies. Microbiome data of single colonies were grouped based on the presence (*mcyA(+)*) or absence (*mcyA*(*-*)) of the microcystin-producing gene *mcyA*. (A) Co-occurrence networks of *mcyA(+)* and *mcyA*(*-*) bacterial communities on ASV level. Nodes were colored according to the respective taxonomic class, except *Microcystis* was colored according to genus. Node size reflects relative abundance. The backgrounds depict example *Microcystis* colonies. (B) Richness of *mcyA(+)* and *mcyA*(*-*) single colony communities. Each dot represents the richness of a single colony in the respective group (*n*(*mcyA*(*-*)) = 17; *n*(*mcyA(+)*) = 12). Box plots show the median (horizontal line), the lower and upper bounds of each box plot indicate the first and third quartiles and whiskers above and below the box plot show 1.5 times the interquartile range. The asterik represents significant difference (*p* < 0.05, Wilcoxon rank sum exact test & Linear regression modelling). (C) Principal Co-ordinates analysis (PCoA) plot on Bray–Curtis dissimilarities of single colony community composition on genus level. The ellipses represent 95% confidence intervals. Color of points, boxes and ellipses correspond to samples with presence (orange) and absence (purple) of *mcyA*-gene.

Based on this observation, we set out to estimate the richness on the genus level for the two groups (Figure 1B). To focus on the effect of MC on the heterotrophic microbiome, cyanobacterial ASVs were excluded prior to this analysis. As already indicated by the findings of the co-presence network, calculation of sample richness confirmed that presence of the *mcyA* gene was indicative for a significant lower richness (*p* < 0.05, Wilcoxon rank sum exact test, LRM). This suggests that MC-producers establish a less diverse and thus a potentially higher specialized microbiome. In order to test the similarity of the phycosphere microbiomes of MC+ and MC- colonies, we used PCoA ordination to visualize Bray-Curtis dissimilarities together with permutational multivariate analysis of variance (PERMANOVA). This demonstrated a significant difference between the chemotypes (PERMANOVA: R^2^ = 0.08434, *p* = 0.013), although it accounted only for 8.4% of the variation, suggesting that MC-production is a minor contributor to the overall variation (Figure 1C). To identify differentially abundant taxa in both groups, we used Linear discriminant analysis effect size analysis (LEfSe analysis) (28). Three taxa were scored in the MC+ colonies: *Tabrizicola*, *Phenylobacterium* and a *Microscillaceae* family member. In the MC- group, two taxa were scored: *Cutibacterium* and *Streptococcus* (Figure S3). Taken together, these findings suggest that the production of MC might support *Microcystis* in their ability to filter heterotrophic bacterial partners in their phycosphere.

### Temporal dynamics of synthetic communities in co-culture with an MC+ and an MC- laboratory strain

To further test the hypothesis that the presence of MC has an influence on the *Microcystis* phycosphere composition we designed a synthetic community experiment with the MC- producing laboratory strain *M. aeruginosa* PCC 7806 (WT) and its MC-deficient Δ*mcyB* mutant. To establish a synthetic consortium, 13 different heterotrophic bacteria were isolated from a *Microcystis* bloom sample of Lake Zernsee. To better represent the natural diversity of *Microcystis* phycosphere microbiomes, we further included eight strains obtained from other phycosphere microbiomes sampled at the University of Jena, Germany or isolated as contaminants in cyanobacterial laboratory cultures (Table S3). Together, the 21 strains covered a diversity of taxa belonging to the classes Alphaproteobacteria, Betaproteobacteria Gammaproteobacteria, Bacteroidia, Actinomycetia and Bacilli broadly resembling natural *Microcystis* phycosphere communities (Figure 2A and Table S3). Final phototroph-heterotroph communities comprised either the PCC 7806 WT, or its MC-deficient Δ*mcyB* mutant together with the defined consortium of 21 strains (Figure 2A and B). Three replicates of axenic cyanobacterial strains and of both consortia were co-cultivated for 28 days. During this time, bacterial growth was monitored by automatic optical density measurement every five minutes at 720 nm. Samples for DNA isolation were collected weekly (Figure 2B). Monitoring of the OD_720_ showed that in both cases (WT and Δ*mcyB* mutant), the co-cultivation condition ultimately reached higher values than the axenic condition (Figure 2C). Measurements of OD_720_ of heterotrophic bacteria alone also displayed absorbance at 720 nm, suggesting that the OD in the co-cultivation is a mixed value of cyanobacterial and heterotrophic bacterial absorbance. However, higher OD values in the co-cultivation could also indicate that the presence of heterotrophic bacteria promoted an overall increased growth rate of *Microcystis* compared to the axenic condition. A mutual influence on the growth dynamics of phototrophs and heterotrophs is indicated by an interesting phenomenon in the community growth curves (Figure 2C): after the initial exponential phase (Day 0 – 9), the cultures transitioned into an intermittent stationary phase, where OD_720_ values visibly receded (Day 10 – 12). Remarkably, the co-cultures advanced into a second exponential growth phase that lasted until the end of the experiment (Day 28). This trend was less pronounced in the axenic WT cultures and not visible in the Δ*mcyB* mutant axenic cultures, suggesting these growth dynamics might be linked to the presence of MC and may be enhanced by heterotrophic bacteria.

**Figure 2.**
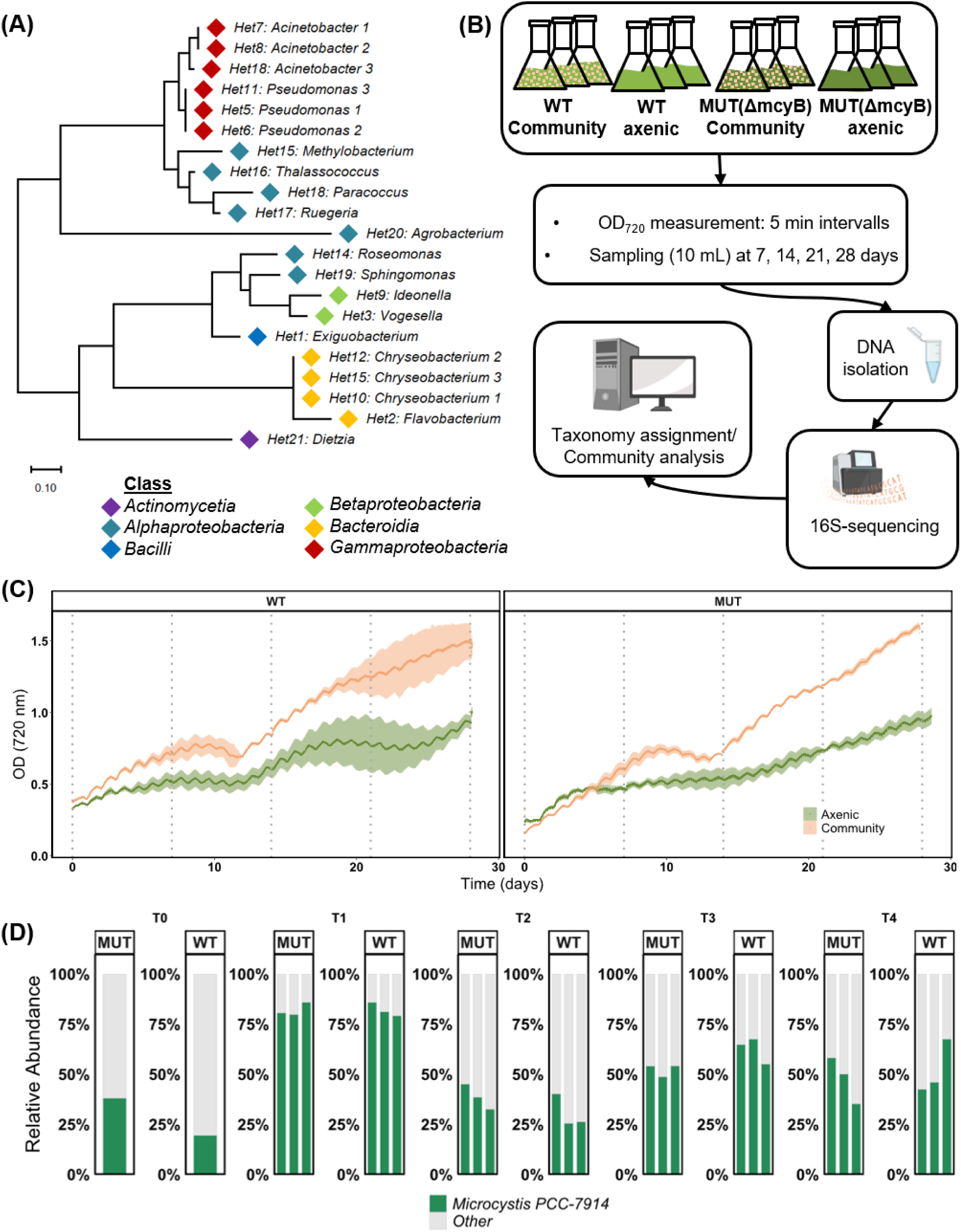
Characteristics of the synthetic community experiment realized through co-cultivation of the MC-producing *Microcystis aeruginosa* PCC 7806 WT or the non-producing Δ*mcyB* mutant together with 21 bacterial isolates. (A) Phylogenetic tree of 21 bacterial isolates used to assemble the heterotrophic bacterial consortium with identified genus label and color-coded by class. Identifiers in front of the colon show ascending enumeration according to the time point each heterotroph was included in the isolate collection. Numbers behind the taxa label indicate separate isolates of the same genus. Detailed descriptions about isolates are shown in Table S3. (B) Illustration of the experimental workflow. Four different conditions were tested: axenic *Microcystis aeruginosa* PCC 7806 (WT axenic), axenic non-producing mutant (Δ*mcyB* axenic), *Microcystis aeruginosa* PCC 7806 WT in co-cultivation with the synthetic consortium (WT Community) and the non-producing mutant in co-cultivation with the synthetic consortium (Δ*mcyB* Community). Three biological replicates were established for each condition (*n* = 3). The cultures were subjected to a day-night-cycle of 15h of constant daylight (55 µmol/m^2^*s) and 7h constant darkness (0 µmol/m^2^*s). Transitions between day and night phases were implemented with linear change of light intensity and the respective target light intensity was reached after 30 min. Automated measurement of optical density (OD) was done at 720 nm throughout the whole course of the experiment every 5 min. Samples were taken weekly over a period of 28 days. Parts of the illustration were created with biorender.com. (C) Development of OD_720_ of *Microcystis aeruginosa* PCC 7806 WT and Δ*mcyB* mutant over time, colored according to experimental condition. Deep-colored lines represent the mean of *n* = 3 (axenic condition) and *n* = 2 (community condition) replicates. Standard deviation is shown as pale ribbons. (D) Relative abundance of *Microcystis* at each sampling timepoint in WT and Δ*mcyB* mutant co-cultivation. Each bar represents a replicate. Reads assigned to *Microcystis* are colored in green, non- *Microcystis* reads are colored in grey.

To estimate the relative abundance of *Microcystis* at different time points we further used 16S- rRNA V3-V4 amplicon sequencing. Taxonomic differentiation of *Microcystis* and non- *Microcystis* amplicons revealed pronounced temporal dynamics in the relative abundance of *Microcystis* during different stages of the co-cultivation experiment. At the beginning of the experiment the initial relative abundance of *Microcystis* was 19,5% and 38,1% in the WT and the Δ*mcyB* mutant strain, respectively (T0, Figure 2D, Table S4). Maximal cyanobacterial relative abundance was observed in both consortia after one week (T1, Figure 2D, Table S4) with 82,0% ± 3,4% in the WT and 82,1% ± 3,3% in the Δ*mcyB* mutant condition, followed by a steep decline after two weeks (WT 30,7% ± 8,3%; Δ*mcyB* mutant 38,7% ± 6,3%). This decline occurred simultaneously with the intermittent stationary phase in the growth curves (Figure 2C), suggesting a potential correlation. From week three onwards, a slight increase in *Microcystis* relative abundance was visible, stabilizing towards the end of the experiment with a final relative abundance of 52% ± 13,6% in the WT and 47,9% ± 11,6% in the Δ*mcyB* mutant, respectively (T4, Figure 2D, Table S4).

Throughout the co-cultivation experiments, the abundance of individual heterotrophic bacteria changed considerably. Although we aimed for similar proportions for all 21 heterotrophs in the inoculum, the genera *Vogesella* (68%) *Chryseobacterium* (19,1%) and *Methylobacterium* (7,8%) were in fact overrepresented at T0 (Figure S4). However, they became less abundant over time and other genera became predominant (Figure 3A, Figure S4). Notably, relative abundance ratios between cyanobacteria and heterotrophs remained stable after two weeks of co-cultivation, especially after T3 (Figure 2D, Figure S4). Assuming that the increase in OD originates primarily from replication, this could indicate that both cyanobacteria and heterotrophic bacteria reproduced in a constant and perhaps even synchronized manner during that period.

**Figure 3.**
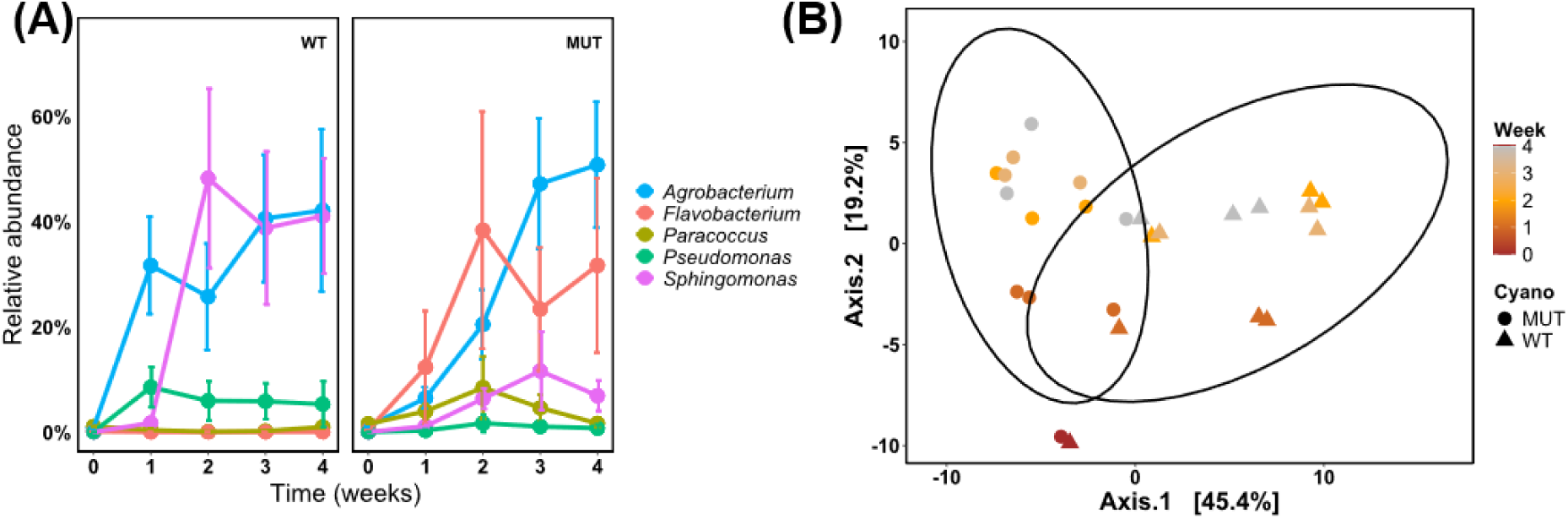
Temporal dynamics of the synthetic community experiment. (A) Relative abundance of selected heterotrophic bacterial taxa over time in the two co- cultivation set-ups with *Microcystis aeruginosa* PCC 7806 and Δ*mcyB* mutant. Taxa were selected based on scoring by LEfSe anaylsis (*p* < 0.05 (Kruskal-Wallis-Test), logLDA cutoff = 3, CSS normalization). Individual taxa are shown in different colors. Filled circles represent mean relative abundance and error bars represent standard deviation (*n* = 3). (B) Principal components analysis (PCA) plot of sample composition on genus level. The first two dimensions are shown. Abundance data were transformed using the centered-log-ratio method. A pseudo count of half of the minimal relative abundance was added to exact zero relative abundance entries in the ASV table. Data points represent samples that are categorized by shape (WT = triangular, MUT (Δ*mcyB*) = circular), and by time (color gradient). The ellipses represent 95% confidence interval. (C) Relative abundance of *Microcystis* and the heterotrophic bacterial consortium over time in the two co-cultivation set-ups. Each bar represents a replicate at the respective time point.

Temporal dynamics in the consortium composition were evaluated by plotting a Principal Components Analysis of the different samples (Figure 3B). At the beginning of the experiments both WT and Δ*mcyB* mutant displayed high community similarity. Already after one week both genotypes seemed to develop their microbiome in different directions. Maximum distance from the initial consortium and between the two genotypes (WT and Δ*mcyB* mutant) was reached after two weeks and remained stable towards the end of the experiment (Figure 3B). The significant differences observed in the overall community composition underline a possible influence of MC on the composition of the microbiome of the *Microcystis* phycosphere.

Using LEfSe analysis, significantly differentially abundant heterotrophs (marker heterotrophs) in each condition were scored (Figure S5). Longitudinal trajectories of the scored bacteria are shown in Figure 3A. Reciprocal abundance development was observed for the *Sphingomonas* and the *Flavobacterium* strains which showed dynamic growth exclusively in the WT and Δ*mcyB* mutant consortium, respectively (Figure 3A). Besides, the *Agrobacterium* strain showed a strong growth in both consortia regardless of the presence of MC.

Taken together, our data suggest that cyanobacteria and heterotrophs took approximately two weeks to build a stable community. MC had a clear impact on the composition of the community and MC-responsiveness was highly strain-specific.

### Metabolic profile and motility characteristics differ between marker heterotrophs

To better understand the influence of MC on the interaction between cyanobacteria and heterotrophs, we focused on three selected strains: *Agrobacterium*, which showed very good growth in *Microcystis* co-cultures independent of MC, and *Sphingomonas* and *Flavobacterium* which thrived exclusively with the WT or the Δ*mcyB* mutant cultures. The three strains were subjected to a metabolic profiling using EcoPlates^TM^, which contain 31 different carbon and nitrogen sources together with a redox dye to indicate substrate specific metabolic activity. To capture the influence of *Microcystis* exudates on strain specific growth and substrate utilization, we additionally used sterile filtered *Microcystis* WT and Δ*mcyB* mutant culture exudates as liquid media for the incubation. Well color development was recorded for 192 h. Only *Agrobacterium* was able to convert substrates without the addition of *Microcystis* exudates, showing broad substrate utilization capabilities including for carbohydrates (e.g. D-cellobiose, D-mannitol, D-lactose), amino acids (e.g. L-asparagine, L-arginine, L-serine) and carboxylic acids (e.g. D-malic acid) but also phosphor containing compounds like glucose-1-phosphate or esters like pyruvic acid methyl ester (Figure 4, Figure S6). The *Microcystis* exudates had only minor impact on substrate utilization. *Sphingomonas*, on the other hand, was only able to show metabolic activity with *Microcystis* exudate supplementation. Both the WT and the Δ*mcyB* mutant exudate enabled substrate utilization on polymeric substrates (α-cyclodextrin, Tween- 40 and Tween-80) and carbohydrates (D-cellobiose and N-acetyl-D-glucosamine). The growth promotion was much more pronounced with WT exudates, where much higher mean well color values were reached compared to the Δ*mcyB* mutant exudates, supporting the hypothesis that the *Sphingomonas* prefers MC+ conditions (Figure 4, Figure S7). The additionally selected *Flavobacterium* was not able to show metabolic activity in any condition, suggesting that other factors are needed for it to be metabolically active (Figure 4, Figure S8). We frequently observed that *Flavobacterium* could not grow in liquid culture, growth was only observed on agar plates. Here, biofilm formation or surface attachment might play an important role to stimulate growth and metabolism.

**Figure 4.**
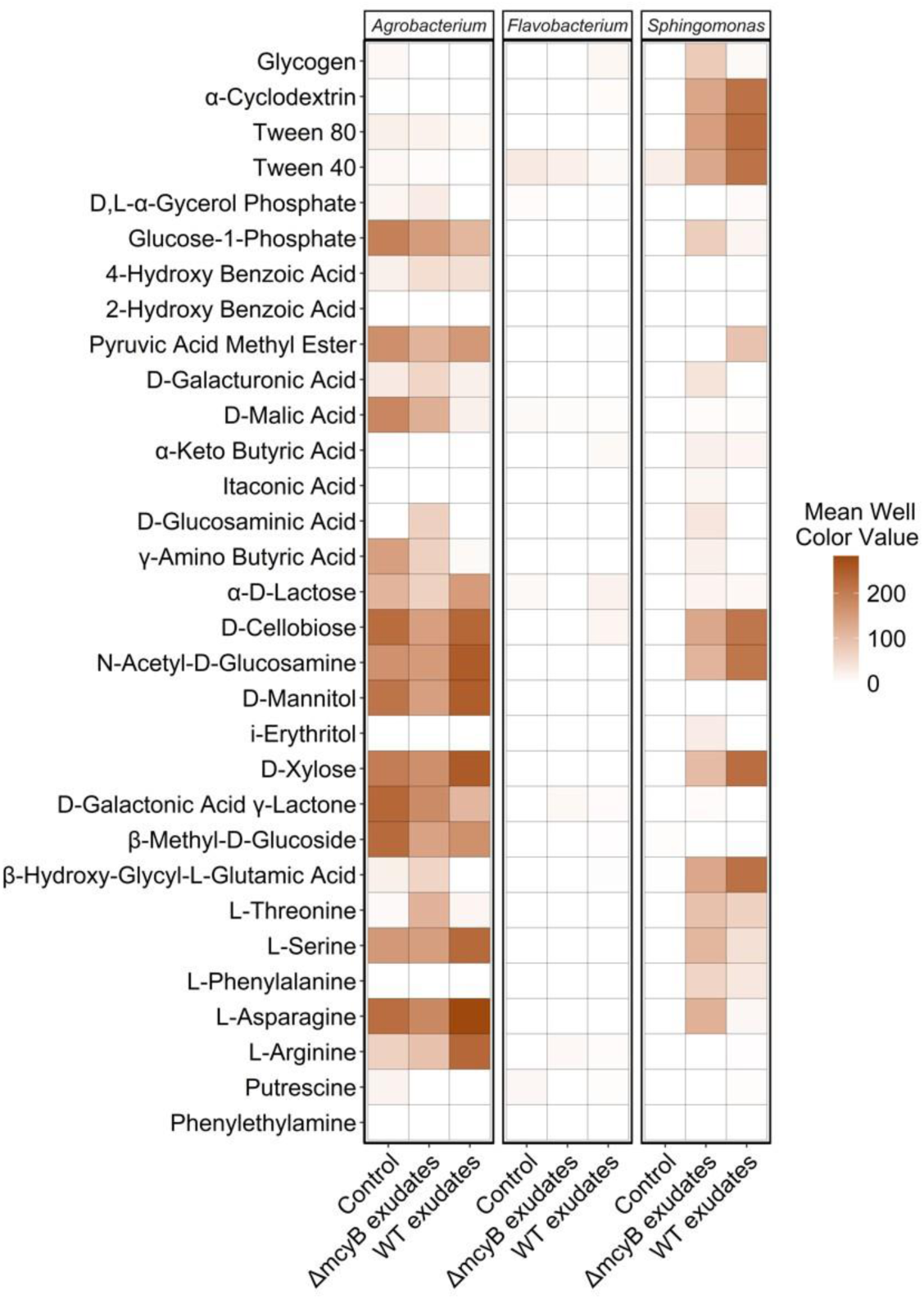
Substrate utilization analysis using EcoPlate^TM^ with and without supplementation of *Microcystis* exudates. Heatmap shows mean well color values of biological replicates (*n* = 3) at the timepoint t = 100 h. Higher well color indicates higher substrate specific metabolic activity of the respective bacterium (*Agrobacterium* sp. UP1*, Flavobacterium* sp. UP2*, Sphingomonas* sp. UP3). Substrate specific metabolic activity was tested in control condition (0.9%-NaCl solution) or in presence of *M. aeruginosa* PCC 7806 WT or Δ*mcyB* mutant exudates. See figures S6-S8 for full 200h time course for individual strains and substrates.

Next, the three selected heterotrophs were studied through whole genome sequencing. This was done using short-read BGI DNA nanoball sequencing (DNBSEQ) and subsequent annotation with bakta (29) and RASTtk (30) analysis. During the course of this analysis, the strains were named *Agrobacterium* sp. UP1, *Flavobacterium* sp. UP2 and *Sphingomonas* sp. UP3. In agreement with the EcoPlate^TM^ analysis, the genome sequence of the *Agrobacterium* sp. UP1 revealed the broadest capabilities to utilize carbohydrate substrates including monosaccharides, carboxylic acids and sugar alcohols, among the three isolates. The genome of *Flavobacterium* sp. UP2, on the other hand, showed the fewest possibilities for utilizing carbohydrates. In particular, we compared the genomic potential to utilize 2-P-glycolate and glycolate, which are among the major dissolved organic carbon species in exudates of phototrophic microorganisms. Genome sequencing revealed that both, *Sphingomonas* sp. UP3 and *Agrobacterium* sp. UP1 encode enzymes required for photorespiration, indicated in the KEGG pathway module “photorespiration” (Figure 5, Table S5). *Flavobacterium* sp. UP2, on the other hand only covered 13.8% of the pathway and did not encode enzymes for the conversion of glycolate or glyoxylate (Table S5). This suggests, that both *Agrobacterium* sp. UP1 and *Sphingomonas* sp. UP3 have the potential to utilize (2-P)-glycolate released by *Microcystis* and either use it as their own substrate (commensal interaction) or to convert it and return glycerate and CO_2_ back to *Microcystis* (mutualistic interaction). As *Microcystis* is not able to produce vitamin B12 we further compared the respective genomic potential of the three strains. Both *Agrobacterium* sp. UP1 and *Sphingomonas* sp. UP3 revealed the ability to produce vitamin B12 according to the KEGG pathway “porphyrin and chlorophyll metabolism”, however they encode different sets of enzymes either for the conversion of precorrin 2 to cob(II)yrinate a,c diamide (*Agrobacterium* sp. UP1) or for a bypass through the amino acids glycine and threonine (*Sphingomonas* sp. UP3) (Table S5). As *Flavobacterium* sp. UP2 does not encode enzymes enabling either conversion it is probably only able to produce vitamin B12 when the required educts are present.

**Figure 5.**
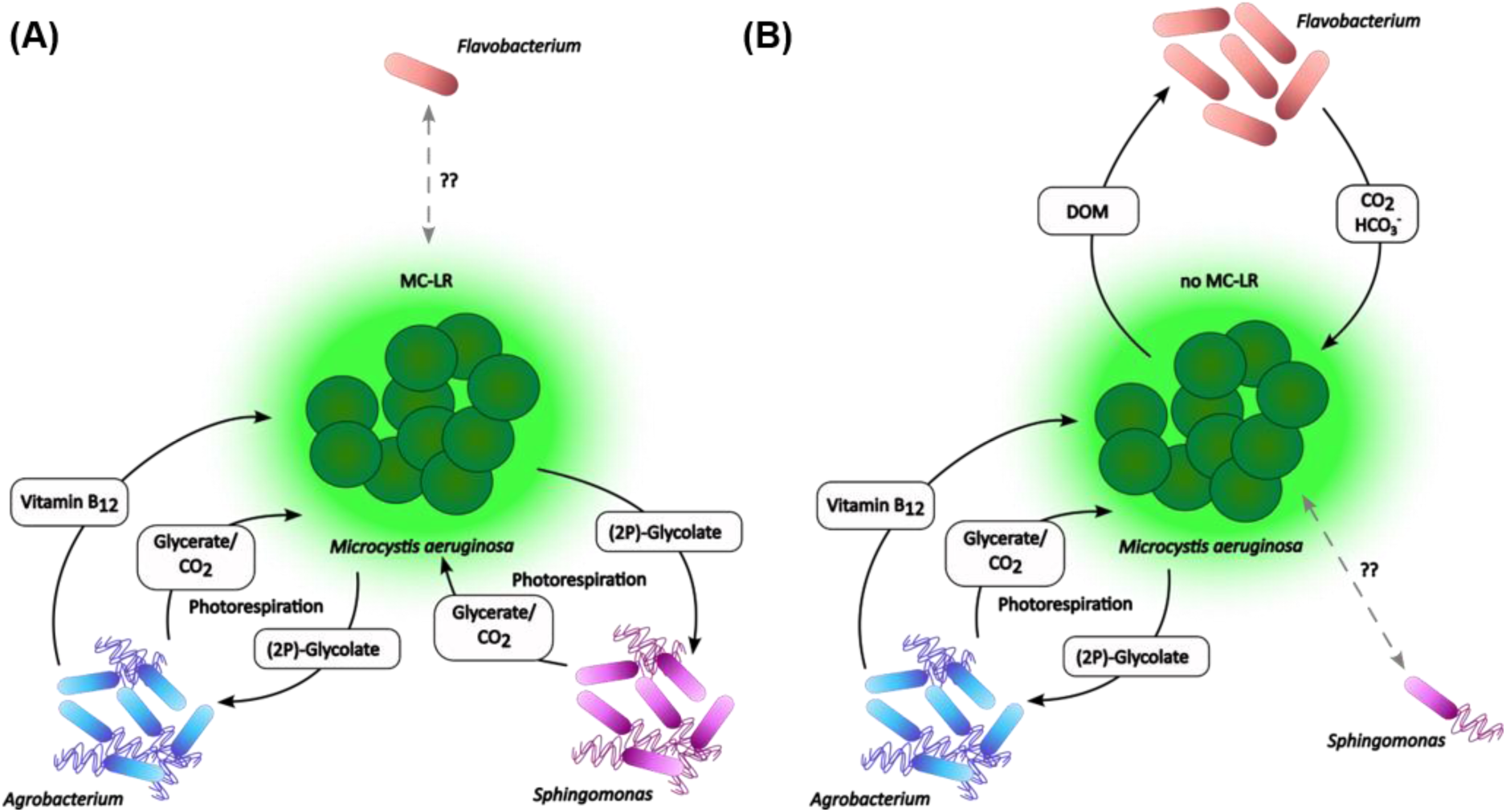
Potential *Microcystis*-heterotroph interactions under MC+ or MC- conditions. Schematic representation of the relationship between *Microcystis* and heterotrophic bacteria summarizing the findings of the SynCom experiment and KEGG pathway analysis based on genome sequencing. Number of heterotrophs in each condition reflects the detected relative abundances in the SynCom experiment (Figure 4A). Putative interaction pathways indicated by arrows are based on the findings of the KEGG pathway analysis of the respective genome. Motility of the heterotrophic bacteria is indicated by flagella. (A) In the presence of MC-LR *Agrobacterium* and *Sphingomonas* reach high relative abundances, whereas *Flavobacterium* is low abundant. Suggested pathways of *Sphingomonas* and *Agrobacterium* are predominant. *Sphingomonas* can perform photorespiration. *Agrobacterium* fulfills a dual role: i) Participate in photorespiration and ii) provide *Microcystis* with vitamin B_12_. (B) In the absence of MC-LR *Agrobacterium* and *Flavobacterium* reach high relative abundances, whereas *Sphingomonas* is low abundant. Suggested pathways of *Agrobacterium* and *Flavobacterium* are predominant. *Flavobacterium* can provide *Microcystis* with CO_2_ and bicarbonate. In return, *Microcystis* provides *Flavobacterium* with dissolved organic matter (DOM). The role of *Agrobacterium* remains equal to (A).

Interestingly, *Flavobacterium* sp. UP2 was found to be uniquely equipped with a bicarbonate transporter (Figure 5). Moreover, we identified differences in the motility potential of the three heterotrophs. Both *Agrobacterium* sp. UP1 and *Sphingomonas* sp. UP3 encode the necessary components of the flagellar biosynthesis apparatus and one or two copies of the chemotaxis regulator CheY, respectively indicating full capabilities for flagellar motility (Figure 5, Table S6 and 8). *Flavobacterium* sp. UP2, again, lacks this ability and only encoded three gliding motility-associated ABC transporter ATP-binding proteins and one putative archaeal flagellar protein (Figure 5, Table S7). Taken together, the EcoPlate^TM^ analysis, *Microcystis* exudate supplementation and genome sequencing indicate that both *Agrobacterium* sp. UP1 and *Sphingomonas* sp. UP3 are ideally equipped for a cross-feeding with *Microcystis*. As *Sphingomonas* sp. UP3 was only able to grow with *Microcystis* exudate supplementation it may more strongly rely on photorespiration than *Agrobacterium* sp. UP1 and may require assistance of *Microcystis* for the degradation of complex polymers like Tween and α-cyclodextrin. In contrast, we were unable to gather any concrete evidence regarding the basis of the interaction between *Microcystis* and *Flavobacterium* sp. UP2 and therefore cannot explain why *Flavobacterium* sp. UP2 showed growth in the MC consortium specifically. We could not find hints for enzymatic MC-degradation via the *mlrA-D* gene cluster (31), since these genes were not identified among the genomes of the three heterotrophs.

## Discussion

In recent years, it has been increasingly recognized that *Microcystis* and its phycosphere bacteria must be regarded as holobiont. There are already numerous indications for a high specificity of the interactions of *Microcystis* and its microbiome and even evidence for a co-evolution of individual traits (6, 7, 12, 17, 32). Yet, the possible role of MC has so far only been touched upon (18). This is mainly due to the fact that the ability to produce MC cannot be clearly assigned to specific *Microcystis* oligotypes but shows a sporadic distribution, at least in some of *Microcystis* genospecies (17, 33). Additionally, most of the studies are still being carried out with bulk *Microcystis* samples, which do not allow a discrimination between toxic and non-toxic genotypes (6, 11). Although *Microcystis* single colonies have already been used to study the *Microcystis* phycosphere composition (12, 17, 27), we performed an independent small-scale single colony analysis at the beginning of this study. Our aim was to focus more exclusively on MC without variation in seasonality, weather conditions, temperature and nutrient concentrations, and to create a basis for the design of the subsequent synthetic community study. This should ultimately lead to a better inter-connection of field and laboratory-based experiments and create the conditions to grasp an integrative functional understanding how MC contributes to the structure of the *Microcystis* phycosphere.

The *Microcystis* single colony analysis initially confirmed many known findings on the composition of the *Microcystis* phycosphere. In particular, we were able to observe many taxa that have already been described for bulk *Microcystis* samples or single colonies. (e.g. *Flavobacterium*, *Roseomonas, Sphingomonas*, *Bradyrhizobium*, *Phenylobacterium*) (18, 27, 10, 34, 35). As described in other studies, we did not observe a *Microcystis* core microbiome community (27). Neither were we able to detect taxa exclusively occurring in MC+ or MC- colonies (17). Yet, MC had a negative impact on community richness and certain taxa preferably occurred in MC+ or MC- colonies. According to the LEfSe analysis, the genera *Tabrizicola*, *Phenylobacterium* and *Microscillaceae* were more prevalent in the MC+ colonies. *Tabrizicola* is a genus belonging to the Rhodobacteraceae family of aerobic anoxygenic phototrophs (AAP). A recent study on metagenome-assembled genomes and metatranscriptomes from *Microcystis* blooms identified AAP bacteria as a particularly relevant group for complementary nutrient recycling in blooms (36). *Phenylobacterium*, was previously described to promote the dominance of MC-producing over non-producing *Microcystis* (18). Therefore, it was not surprising to find this genus associated, especially in MC+ colonies. The genera *Cutibacterium* and *Streptococcus*, on the other hand, which were more strongly associated with MC- colonies, are rather atypical for *Microcystis* microbiomes. Their prevalence in MC- colonies may reflect the overall greater species richness in MC- colonies. Both genera are known to be present in the human skin microbiomes and their occurrence may be due to the fact that Lake Zernsee is a very popular swimming water. However, this might also indicate that MC itself or the MC-dependent phenotypic differences of *Microcystis* act as a selective filter in the *Microcystis* phycosphere. Overall, our *Microcystis* field microbiome analysis suggests that MC may indeed act as chemical mediator contributing to the specificity of *Microcystis* microbiome interactions. However, since most of the MC+ preferring genera such as *Tabrizicola*, *Phenylobacterium* and *Microscilla* occur in co-occurrence networks of both colony types and only the individual ASVs differ, functional traits that are not part of the core genome of these genera are probably relevant for the MC preference. According to our study, genus assignment is therefore not a good indicator for MC+ or MC- colony specificity.

Individual *Microcystis* colonies differ in a large number of functional traits such as their sheath properties and inorganic carbon adaptation, which poses a great challenge to capture the distinct influence of MC. The synthetic community experiment with the MC-producing strain PCC 7806 and its Δ*mcyB* mutant was therefore designed to reduce the complexity, both with regard to the functional traits and to the number of involved taxa. In this reductionist approach, the strong mutual influence of the partners in our phototroph-heterotroph consortia is already reflected in the growth curves of the consortia, which indicate a real tug-of-war between the partners (Figure 2, Figure S4). In the ultimately stable consortium, however, the interactions appear to be mainly mutualistic, as the three heterotrophic isolates and also *Microcystis* itself grew during this period. A very clear influence of MC could be demonstrated and even strains could be identified that were exclusively thriving with one or the other genotype. This strong influence of a secondary metabolite on the community composition is quite remarkable. A recent study on the impact of antibiotic production in *Bacillus* strains on the composition of a community of 13 distinct genera of soil bacteria revealed only marginal differences when either the *Bacillus* WT strain or mutants defective in surfactin, plipastatin or bacillaene biosynthesis were co-cultured despite the strong antibacterial effects of these metabolites (37).

For the bacteria of the *Microcystis* phycosphere, a major role through complementary nutrient recycling is being discussed (6, 36). We therefore examined the extent to which cross-feeding could contribute to a mutualistic relationship between the partners for the three selected indicator organisms *Agrobacterium*, *Sphingomonas* and *Flavobacterium*. Based on whole genome sequencing and KEGG pathway analysis, we observed a complementary metabolic potential especially for *Agrobacterium* and *Sphingomonas*. In particular, their ability to perform photorespiration, and the production of vitamin B12 can complement *Microcystis* metabolism. Recently, it was shown that an *Agrobacterium* strain can efficiently compensate for the inorganic carbon limitation of the *Nostoc* strain *N. punctiforme* PCC 73102 (38). Since many *Microcystis* strains have a weak carbon concentrating mechanism (CCM) (39) and inorganic carbon is a limited resource in blooms (40), we also assume that a major contribution of the selected heterotrophic bacteria is the supply of respiratory CO_2_ to *Microcystis* (Figure 5).

The fact that *Sphingomonas* grew better with WT exudates suggests that MC plays an active role especially in cross-feeding between *Microcystis* and *Sphingomonas*. We hypothesize that *Sphingomona*s is more dependent on photorespiration than *Agrobacterium*. It has been known for some time that MC has a very prominent effect on the inorganic carbon metabolism of *Microcystis* and, in particular, influences the accumulation of RubisCO products (24, 25). It is therefore possible that MC+ conditions could favor nutrient recycling and in turn the interaction with *Sphingomonas*. Our analysis showed that in the presence of MC, *Sphingomonas* was able to utilize more carbon sources, including complex substrates such as Tween-40, 80 and α- cyclodextrin. This, in turn, suggests that *Microcystis* could potentially stimulate or be involved in the degradation of these complex substrates. However, nothing is known yet about corresponding enzyme activities in *Microcystis*. *Sphingomonas* is one of the bacterial genera frequently partnering with *Microcystis* and is mainly studied in connection with the degradation of MC. However, the strain selected in this study is not equipped with the known MC degradation enzymes *mlrA-D*. When looking at the co-occurrence networks of MC+ and MC- colonies, it is noticeable that one individual *Sphingomonas* ASV is linked with MC- colonies suggesting that also for this genus functional traits which are not part of the genus’ core genome are decisive for the interaction. Indeed, pathways such as MC degradation, photorespiration and vitamin B12 production are only sporadically encoded in *Sphingomonas* genomes and could influence the interaction in different ways (Table S6). In agreement with this hypothesis, Berg *et al*. have already shown that different *Sphingomonas* isolates had diverse and even opposite outcomes on growth of *Microcystis* using a cultivation-based approach (10).

A similar observation was also made for *Flavobacteria* which are also frequently co-associated with *Microcystis* blooms. Here too, either growth-promoting or growth-inhibiting influences on *Microcystis* were detected (10) and *Flavobacteria*-ASVs also occur in both MC+ and MC- colonies in our co-occurrence networks (Figure 1A). In our synthetic community study, the selected *Flavobacterium* isolate showed a clear preference for the Δ*mcyB* mutant strain. However, neither the EcoPlate^TM^ analyses nor the genome sequence provide information on traits that could be pivotal for the interaction with *Microcystis* or explain the Δ*mcyB* preference. It is possible that the interaction of the partners in this case requires a physical interaction that cannot be reproduced by the addition of exudates. As PCC 7806 and its Δ*mcyB* mutant are known to differ in their cell surface characteristics, for example in the expression of the cell surface glycoprotein MrpC, a preferential interaction with one of the genotypes could also be rooted in alterations of cell-cell interaction characteristics.

In summary, both the field experiment and the synthetic community experiment suggest that MC has an influence on the structure of the phycosphere microbiome of *Microcystis*. This effect was more pronounced in the synthetic community analysis. This seems logical, as in the more complex field study the influence of MC is interfering with other factors such as the morphotype and sheath characteristics or the different adaptation of *Microcystis* strains to different inorganic carbon conditions. The subsequent functional study was able to show that MC probably influences cross-feeding between the strains through its influence on inorganic carbon metabolism and/or by promoting the degradation of complex substrates. The study thus further emphasizes previous findings on the role of *Microcystis* phycosphere bacteria in complementary nutrient recycling and highlights the great importance of strain-specific traits for the nature of interactions and the response to MC production.

## Materials and methods

### Single colony isolation

Single *Microcystis* colonies were sampled during a bloom event on July 14^th^, 2021 on three locations along the lake Zernsee (Mühlendamm: 52.417028741058964, 12.935628335139363; Gohlwerder: 52.428188547155465, 12.936549043106284; Schwarzer Weg: 52.425179744399784, 12.93285196111643). *Microcystis* colonies were sampled from the water surface, using a plankton net and transferred into 50 mL Falcon tubes. During transportation the samples were kept in the dark at 4°C. Colony isolation was done under the stereo microscope Stemi 305 (Zeiss, Oberkochen, Germany) following the “Isolation Using Micropipettes” method described by Kurmayer, R. *et al.* (41) within 36 h after sampling. Images of single colonies were acquired with the Zeiss Axiocam microscope camera. Single colonies were stored in 10 µL sterile H_2_O at -20°C until DNA extraction.

### Heterotrophic strain isolation and maintenance

Heterotrophic bacteria were isolated from lake Zernsee water samples (see Single colony isolation) and non-axenic cyanobacterial lab strains. Briefly, one drop of lake water containing 1 – 2 *Microcystis* colonies was streaked on CYA Agar (10) or R2A agar plates (Carl Roth, Karlsruhe, Germany) and incubated at 25°C in the dark. After two days, colonies were picked and streaked again. This procedure was repeated until single colonies with unique bacterial species could be obtained, based on visual inspection of colony morphology and color.

From non-axenic cyanobacterial lab strains, 10 µL were streaked on R2A agar plates. Bacteria isolation procedure was done identically to the lake water samples.

Heterotrophic cultures were maintained on R2A agar plates at RT (20 – 25°C).

### Heterotrophic bacterial strain identification

Heterotrophic bacterial isolates were identified using Sanger Sequencing of the 16S-rRNA phylogenetic marker gene. A single colony was picked from an R2A agar plate and resuspended in 20 µL of sterile MilliQ water. 10 µL were used immediately to inoculate a fresh R2A agar plate to ensure that the taxon identified after sequencing and the subsequent bacterial cultures are of the same origin. For the amplification of the 16S-rRNA gene the following primers were used: 16S_27_FW (5’-AGAGTTTGATCCTGGCTCAG-3’) and 16S_1492_RV (5’-GGTTACCTTGTTACGACTT-3’) (42). PCR with Phusion™ High Fidelity DNA polymerase (Thermo Fisher Scientific, Waltham, MA, USA) was carried out using 1.5 µL of the bacterial suspension as template. PCR reaction was run according to the manufacturers protocol with the following changes: Initial denaturation step was extended to 10 min to break down the bacterial cells. Successful amplification of the product was confirmed by running 5 µL of the PCR reaction on a 1% agarose gel using SYBRsafe stain (Invitrogen, Waltham, MA, USA). The PCR product was purified using the GeneJET PCR purification kit (Thermo Fisher Scientific). Final PCR product concentration and quality was measured using the NanoDropOne (Thermo Fisher Scientific). Sanger sequencing of purified amplicons was done by LGC genomics (LGC Ltd, Teddington, England), using the same primer pair. Only 16S-rRNA sequences with a minimal length of 415 bp and a quality score ≥ 40 were used in a BLAST nucleotide search. The extract_regions_16s tool (https://github.com/AlessioMilanese/extract_regions_16s) was used to obtain the V6 regions. V6 regions were copied to a FASTA file and used to construct a Maximum-likelihood phylogenetic tree with bootstrapping of 100 iterations using the Tamura 3-parameter model in the MEGA X software (version 11.0.13).

### Culture maintenance

Maintenance cultures of cyanobacterial strain *Microcystis aeruginosa* PCC 7806 wild type (WT) were grown in BG11 medium (43). For the MC-LR deficient Δ*mcyB* mutant 5 µg/mL Chloramphenicol were added to the BG11 medium. Both strains were cultivated on the benchtop with natural day-night cycles at 20 – 30 °C. Culture flasks were shaken once a day. To assure exponential growth, cultures were passaged when the OD_750_ of cultures reached 1.0 and OD_750_ of fresh cultures was adjusted to approximately 0.2. Axenicity of cultures was monitored regularly by streaking a small sample on R2A agar plates and via microscopy.

Heterotrophic bacterial cultures were grown on R2A agar plates, except strains ENV4 and ENV3 were grown on Marine Agar (detailed information in Table S3), in the dark at 20 – 30°C. Passaging of heterotrophic cultures was done monthly. For the experiments, only bacteria were used that were passaged not more than 5 times.

### Co-cultivation experiment

For the analysis of synthetic communities, a co-cultivation experiment was conducted, in which *M. aeruginosa* PCC 7806 WT and Δ*mcyB* mutant were supplied with a consortium of 21 heterotrophic bacterial isolates (Figure 2A, Table S3) and compared with the growth of axenic cultures. The experiment was run in triplicates. The resulting co-culture should consist of 50% *Microcystis* cells and 50% heterotrophic bacterial cells whereas the axenic cultures should consist of 100% *Microcystis* cells. For the experiment set up, pre-cultures of WT and Δ*mcyB* mutant were centrifuged (4700*g, 10 min, RT) and resuspended in fresh BG11 medium to a final OD_750_ of 0.2, which corresponded to a cell number of 6*10^6^ cells/mL. Pre-cultures of heterotrophs were grown in their respective liquid media for 2 – 4 days. Heterotrophic bacteria were harvested by centrifugation (4700*g, 10 min, RT) and resuspended in fresh BG11 medium. Bacterial cell suspensions were diluted so that the final amount of each heterotroph isolate should have a cell number of 1/21^th^ of the *Microcystis* cell number (∼3*10^5^). The experiment was run over a time course of 4 weeks and samples for DNA extraction were taken weekly. The sample volume was 10 mL. For the incubation, the MultiCultivator OD-1000 (Photon Systems Instruments, Drásov, Czech Republic) was used, which allowed for accurate light and temperature control. The cultivation was done in BG11 medium at 25°C, with a Day-Night-Rhythm (15.5 h light (55 µE), 30 min linear light reduction phase to 0 µE, 7.5 h of darkness (0 µE) followed by a linear light increasement phase from 0 to 55 µE in 30 min repeated 28 times). Growth monitoring was done by the measuring the OD_720_ and OD_600_ automatically every 5 min. Axenicity of cyanobacterial monocultures was confirmed by 16S- rRNA sequencing.

### DNA isolation

DNA extraction from the single colonies was done using the ChargeSwitch® gDNA Mini Bacteria Kit, as previously described by Pérez-Carrascal, *et al*. (17). In brief, cells of single colony samples were lysed in two steps: First, Resuspension Buffer mix containing RNase A and lysozyme were added and samples were incubated at 37° for 20 min. Second, 500 µL Lysis Buffer/Proteinase K mix were added, followed by another incubation step (1h at 80°C). Subsequently, DNA was purified using 40 µL ChargeSwitch® Magnetic Beads per sample. Finally, DNA was eluted in 50 µL Elution buffer by incubating the samples at 65°C for 10 min. DNA extraction from synthetic communities was performed using the DNeasy Blood&Tissue kit (Qiagen, Hilden, Germany). First, cells from 10 mL sample volume were harvested (4700xg, 10 min, 4°C), resuspended in 900 µL ATL buffer and added with 0.5 g glass beads (0,5 mm) and 0.5 g glass beads (0,1mm), followed by a beating step at maximum speed on a vortex for 45 seconds. Cells were then immediately incubated at 56°C for 30 min. Once the incubation was finished, the bead beating step was repeated. Then, 100 µL Proteinase K were added and cells were incubated at 56° (2h). Beads were collected at the bottom of the tube via centrifugation (2 min, 1000xg, 4°C), bead-free supernatant was transferred to a fresh tube and centrifuged again (1 min, max speed, RT). Subsequently, 650 µL of the supernatant were mixed with 650 µL of non-denatured ethanol and transferred to a mini spin column. DNA purification was done according to the manufacturers protocol with a final elution step in 150 µL DNA-free water. All steps were performed under sterile conditions.

### Processing of the single colonies data set

The chemotype of 50 single *Microcystis* colonies was assessed using specific PCR based detection. Single colonies were classified into *mcyA(+)* and *mcyA*(*-*) based on the presence/absence of an NRPS gene from the MC biosynthetic gene cluster *mcyA*. For that the *mcyA*-condensation domain primers *mcyA*-Cd-1R (5′-AAAAGTGTTTTATTAGCGGCTCAT- 3′) and *mcyA*-Cd-1F (5′-AAAATTAAAAGCCGTATCAAA-3′) were used (44). PCR with Phusion™ High Fidelity DNA polymerase (Thermo Fisher Scientific) was carried out using 1 µL of the extracted DNA as template. PCR reaction was run according to the manufacturers protocol. Amplicons of the expected band size (291 bp) were observed on a 3% agarose gel using SYBRsafe stain (Invitrogen). 13 colonies that exhibit a strong PCR band signal were classified as *mcyA(+)*. 18 colonies that showed no PCR band signal were classified as *mcyA*(*-*) colonies (Figure S2). Twenty-one colonies with weak PCR signal were considered “mixed” type (due to incomplete separation of MC-producing and non-producing genotypes). These were not classified (NC) and excluded from further analysis. Additionally, one toxic and one nontoxic colony sample (Sample 19 and Sample 20, respectively) were excluded from the analysis due to atypical high abundances of Vampirivibrionia Class, that was not observed in any other sample. The remaining dataset of 12 *mcyA(+)* and 17 *mcyA*(*-*) colonies were used for the subsequent analysis.

### 16S-rRNA sequencing and microbial community analysis

Microbiome analysis was performed based on the sequencing of the 16S-rRNA V3-V4 region, using the universal primer pair 806R (5’-GGACTACHVGGGTWTCTAAT-3’) and 341F (5’- ACTCCTACGGGAGGCAGCAG-3’). Single colony sequencing was done on the Illumina MiSeq PE300 platform and resulted in a total number 383,582 quality checked reads (BGI Genomics Co., Ltd). For assessment of sequencing accuracy and quality, a mock community was included.

For the synthetic community experiment the DNBSEQ platform was used, which yielded 906,902 quality checked reads (BGI Genomics Co., Ltd).

Raw reads were trimmed using cutadapt version 2.10 (parameter settings: --quality-cutoff 21 \ --minimum-length 213) (45) and quality checked using FASTQC and MULTIQC (46, 47). High quality reads were merged using the DADA2 pipeline (48). Statistical analysis of sequence data was done in RStudio (2024.04.2+764 Chocolate Cosmos) with R version 4.3.2, using the microViz package (https://david-barnett.github.io/microViz/) (49). Relative abundances were calculated using the “compositional” approach. Richness was calculated using the exp_shannon function. LEfSe analysis was done with the microbiomeMarker package with the following parameters: *p* < 0.05 (Kruskal-Wallis-Test), logLDA cutoff = 3, CSS normalization (50). Microbial association networks were constructed using CoNet with its default parameters and visualized in Cytoscape version 3.10.2 (51).

### Substrate utilization test with EcoPlates^TM^

Utilization of specific substrates by the three heterotrophic bacterial strains *Agrobacterium, Flavobacterium, Sphingomonas* was done using EcoPlate^TM^ (Biolog Inc., Hayward, CA, USA). EcoPlates^TM^ are 96-well-plates, which are equipped with a triplicate of 31 carbon sources. Bacterial strains were grown on R2A agar plates at 25°C until sufficient biomass was yielded. In order to test the influence of cyanobacterial exudates on bacterial growth and substrate utilization, *M. aeruginosa* PCC 7806 (WT) and *ΔmcyB* mutant were grown in 200 mL BG11 medium for 8 weeks without medium exchange to enrich cyanobacterial exudates in the medium. The cultures were centrifuged (4700 rpm, 10 min, RT) and subsequently the exudates containing supernatant was collected, sterile filtered (0,2 µm) and stored at 4°C until further use. For the substrate screening experiment, bacterial biomass was scraped off the agar plates and resuspended either in sterile filtered WT/ Δ*mcyB* mutant exudates or 0.9%-NaCl solution (according to the manufacturers protocol). Turbidity of the suspension was adjusted to 60%. For each bacterial strain, one plate was prepared for each condition, resulting in a total of 9 plates. Plates were incubated at 25°C for 192 h in the OmniLog 50 incubator using the OmniLog Data Collection Software 3.0. After the incubation was finished, data were transferred to in the Data Analysis Software 1.7 and exported. Heatmap and kinetic plots were created in RStudio (2024.04.2+764 Chocolate Cosmos) with R version 4.3.2 using the ggplot2 package.

### Genome sequencing of selected heterotrophs

For whole genome sequencing of the three selected heterotrophic bacterial strains (*Agrobacterium, Flavobacterium, Sphingomonas*), bacteria were freshly taken from cryo- preserved culture and grown on R2A agar plates until sufficient biomass was yielded. The cells were harvested and diluted in ATL buffer (Qiagen). DNA was extracted according to the protocol for the synthetic communities mentioned above. High quality genomic DNA was sequenced, using DNBSEQ method (BGI Tech Solutions Co., Hong Kong, China). Raw reads were quality checked using FASTQC (46). Genomes were assembled using shovill (https://github.com/tseemann/shovill) and assemblies were inspected and evaluated using Bandage (52) and Quast (53). Assembled genomes were annotated using the bakta pipeline (29) and the annotation server tool RAST following the RASTtk annotation scheme with default settings (30). Subsequent KEGG comparison and genome browsing was done with the SEED Browser (54).

## Data deposition

The raw sequence reads obtained in this study have been deposited in the Sequencing Read Archive under BioProject number PRJNA1148368. Remaining data and scripts can be obtained at https://gitup.uni-potsdam.de/ag_mibi/NSMCA.

## Acknowledgements

This work was supported by a grant of the German Research Foundation (DFG, Project-ID 239748522- SFB 1127) to E.D. We thank Juniorprof. Dr. Julie Zedler from University of Jena, Germany, and her group for providing heterotrophic bacterial strains ENV2, ENV3, ENV4, ENV5 and ENV6.

## Author contributions

E.D. designed the work, R.G. performed experiments, R.G., M.H., J.E.T., D.K.R., A.A. contributed to data analysis and interpretation, R.G. and E.D. wrote the manuscript with contributions from all authors.

