## Supplementary Tables S1-S4, Supplementary Figures S1-S8 for "Microcystin shapes the *Microcystis* phycosphere through community filtering and by influencing cross-feeding interactions"

**Table S1**: ASV table of 29 single *Microcystis* colonies. ASVs shown were assigned to the genus *Microcystis*-PCC7941 with more than 1000 reads in at least one sample. Green = colonies with dominant ASV2, blue = colonies with dominant ASV3, red = colonies with no dominant ASV, yellow = colonies with dominant ASV1.

| Sample No | Toxicity | ASV1 | ASV2 | ASV3 | ASV4 |
|---|---|---|---|---|---|
| **10** | **T** | 0 | 12613 | 0 | 0 |
| **13** | **T** | 0 | 11037 | 0 | 0 |
| **15** | **T** | 0 | 7058 | 0 | 0 |
| **17** | **T** | 0 | 8894 | 0 | 0 |
| **23** | **NT** | 0 | 1251 | 0 | 0 |
| **25** | **NT** | 0 | 9429 | 0 | 0 |
| **26** | **T** | 0 | 10924 | 0 | 0 |
| **27** | **NT** | 0 | 14687 | 0 | 0 |
| **30** | **NT** | 0 | 10026 | 0 | 0 |
| **31** | **T** | 0 | 9396 | 0 | 0 |
| **36** | **NT** | 0 | 10842 | 0 | 0 |
| **44** | **NT** | 0 | 9942 | 0 | 0 |
| **21** | **NT** | 745 | 0 | 12939 | 0 |
| **7** | **T** | 2681 | 6694 | 0 | 0 |
| **47** | **NT** | 3676 | 0 | 11808 | 0 |
| **38** | **NT** | 5423 | 2735 | 0 | 0 |
| **45** | **NT** | 7766 | 2688 | 0 | 361 |
| **8** | **T** | 8132 | 9007 | 0 | 0 |
| **43** | **NT** | 9638 | 1181 | 0 | 1227 |
| **24** | **NT** | 8254 | 0 | 0 | 0 |
| **11** | **T** | 9257 | 0 | 0 | 0 |
| **48** | **NT** | 9498 | 814 | 0 | 0 |
| **32** | **NT** | 10254 | 0 | 0 | 0 |
| **6** | **NT** | 12973 | 0 | 0 | 0 |
| **28** | **NT** | 13163 | 0 | 0 | 0 |
| **1** | **T** | 15232 | 0 | 0 | 0 |
| **2** | **T** | 15572 | 0 | 0 | 503 |
| **39** | **NT** | 6186 | 0 | 0 | 0 |
| **16** | **T** | 7616 | 0 | 0 | 389 |

**Table S2**: Relative presence of ten most prevalent taxa in 29 single *Microcystis* colonies. Taxonomic resolution is the genus level.

| Genus | Presence in percent of colonies |
|---|---|
| ***Roseomonas*** | 75,9% |
| ***Microscillaceae Family*** | 75,9% |
| ***Vibrio*** | 72,4% |
| ***Cutibacterium*** | 69,0% |
| ***Pelomonas*** | 69,0% |
| ***Tabrizicola*** | 69,0% |
| ***Phenylobacterium*** | 62,1% |
| ***Flavobacterium*** | 62,1% |
| ***Staphylococcus*** | 55,2% |
| ***UKL13-1*** | 48,3% |

**Table S3**: Overview of heterotrophic bacterial strains and media used for the assembly of the heterotrophic consortium to conduct the co-cultivation experiments.

| Het ID | Genus | Source | Culture media |
|---|---|---|---|
| Het 1 | *Exiguobacterium* | Environment: Havel, Gohlwerder | R2A (DSMZ No 830) |
| Het 2 | *Flavobacterium* | Environment: Havel, Gohlwerder | R2A (DSMZ No 830) |
| Het 3 | *Vogesella* | Environment: Havel, Gohlwerder | R2A (DSMZ No 830) |
| Het 4 | *Pseudomonas* | Environment: Havel, Schwarzer Weg | R2A (DSMZ No 830) |
| Het 5 | *Pseudomonas* | Environment: Havel, Schwarzer Weg | R2A (DSMZ No 830) |
| Het 6 | *Acinetobacter* | Environment: Havel, Schwarzer Weg | R2A (DSMZ No 830) |
| Het 7 | *Acinetobacter* | Environment: Havel, Schwarzer Weg | R2A (DSMZ No 830) |
| Het 8 | *Ideonella* | Environment: Havel, Mühlendamm | R2A (DSMZ No 830) |
| Het 9 | *Chryseobacterium* | Environment: Havel | R2A (DSMZ No 830) |
| Het 10 | *Pseudomonas* | Environment: Havel | R2A (DSMZ No 830) |
| Het 11 | *Chryseobacterium* | Environment: Havel | R2A (DSMZ No 830) |
| Het 12 | *Chryseobacterium* | Environment: Havel | R2A (DSMZ No 830) |
| Het 13 | *Acinetobacter* | Environment: Havel | R2A (DSMZ No 830) |
| Het 14 | *Roseomonas sp.* | Provided by AG Zedler, Friedrich-Schiller Universität Jena | R2A (DSMZ No 830) |
| Het 15 | *Methylobacterium* | Provided by AG Zedler, Friedrich-Schiller Universität Jena | R2A (DSMZ No 830) |
| Het 16 | *Thalassococcus* | Provided by AG Zedler, Friedrich-Schiller Universität Jena | Marine Medium (DSMZ No 123) |
| Het 17 | *Ruegeria* | Provided by AG Zedler, Friedrich-Schiller Universität Jena | Marine Medium (DSMZ No 123) |
| Het 18 | *Paracoccus sp* | Provided by AG Zedler, Friedrich-Schiller Universität Jena | R2A (DSMZ No 830) |
| Het 19 | *Sphingomonas* | Lab strain: non-axenic *Synechocysti*s PCC 6803 (RMA mutant) | R2A (DSMZ No 830) |
| Het 20 | *Agrobacterium* | Lab strain: non-axenic *Synechocystis* PCC 6803 (RMA mutant) | R2A (DSMZ No 830) |
| Het 21 | *Dietzia* | Lab strain: non-axenic *Nostoc punctiforme* PCC 73102 | R2A (DSMZ No 830) |

**Table S4**: Mean relative abundances of *Microcystis* in the synthetic communities at the respective timepoint (*n* = 3).

| | ***M. aeruginos*a WT** | | ***ΔmcyB* MUT** | |
|---|---|---|---|---|
| | **Mean relative abundance** | **Standard deviation** | **Mean relative abundance** | **Standard deviation** |
| **T0** | 19,5% | NA | 38,1% | NA |
| **T1** | 82,0% | 3,4% | 82,1% | 3,3% |
| **T2** | 30,7% | 8,3% | 38,7% | 6,3% |
| **T3** | 62,5% | 6,5% | 52,3% | 3,1% |
| **T4** | 52,0% | 13,6% | 47,8% | 11,6% |

**Supplementary figures**

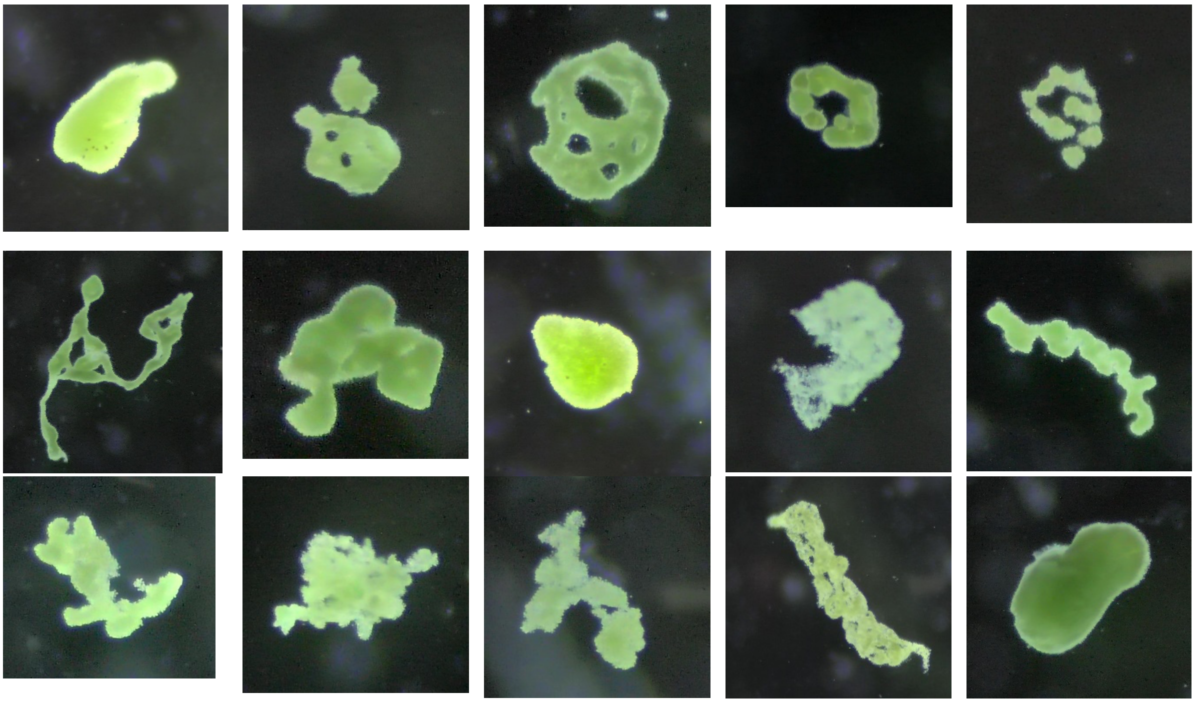

**Figure S1**: Gallery of representative *Microcystis* colonies collected from the Havel river in the Potsdam area (Tiefer See, 52.402987997097476, 13.079468904213506), multiple sampling time points during July and August 2023. Images acquired with Zeiss Stemi 305, equipped with external microscope camera, 10x magnification.

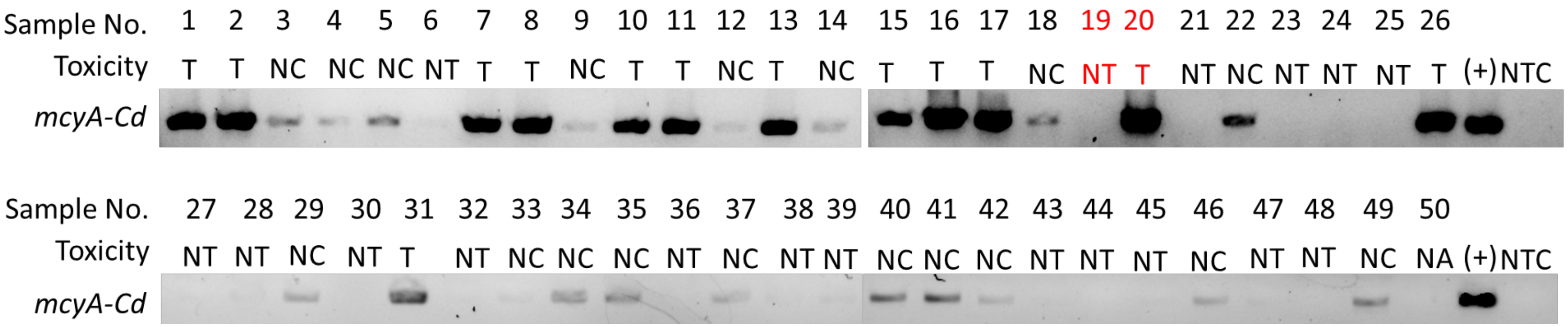

<u>Legend</u>

| | | |
|---|---|---|
| T | – | MC-producing (toxic) |
| NT | – | non-producing (non-toxic) |
| NC | – | excluded from analysis (not classified) |
| Red | – | excluded from analysis (unregular abundance of Vampirivibrionia class) |
| (+) | – | positive control (*M. aeruginosa* PCC 7806 gDNA) |
| NTC | – | negative control (no template control) |

**Figure S2**: Agarose gel electrophoresis of *mcyA-Cd* fragment amplicons. DNA isolated from single *Microcystis* colonies was used as template. Band size of about 300 bp was confirmed with the GeneRuler Low Range DNA Ladder (Thermo Scientific). Colony enumeration and classification into toxic and non-toxic colonies is indicated by letters above each band.

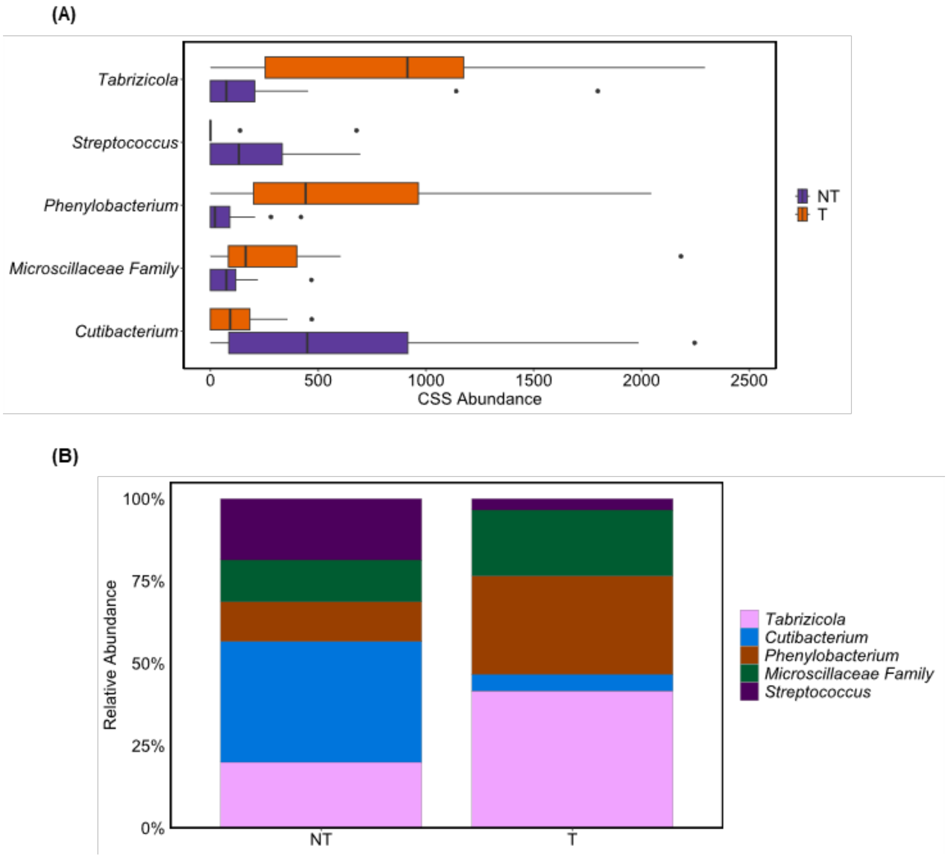

**Figure S3**: Differential abundance analysis of taxa on genus level of single *Microcystis* colonies. (A) Boxplot of CSS abundance of taxa scored by LEfSe analysis of *Microcystis* single colonies. (B) Mean relative abundances of taxa scored by LEfSe analysis. NT = non-MC-producing (MC-) (*n* = 17), T = MC-producing (MC+) (*n* = 12).

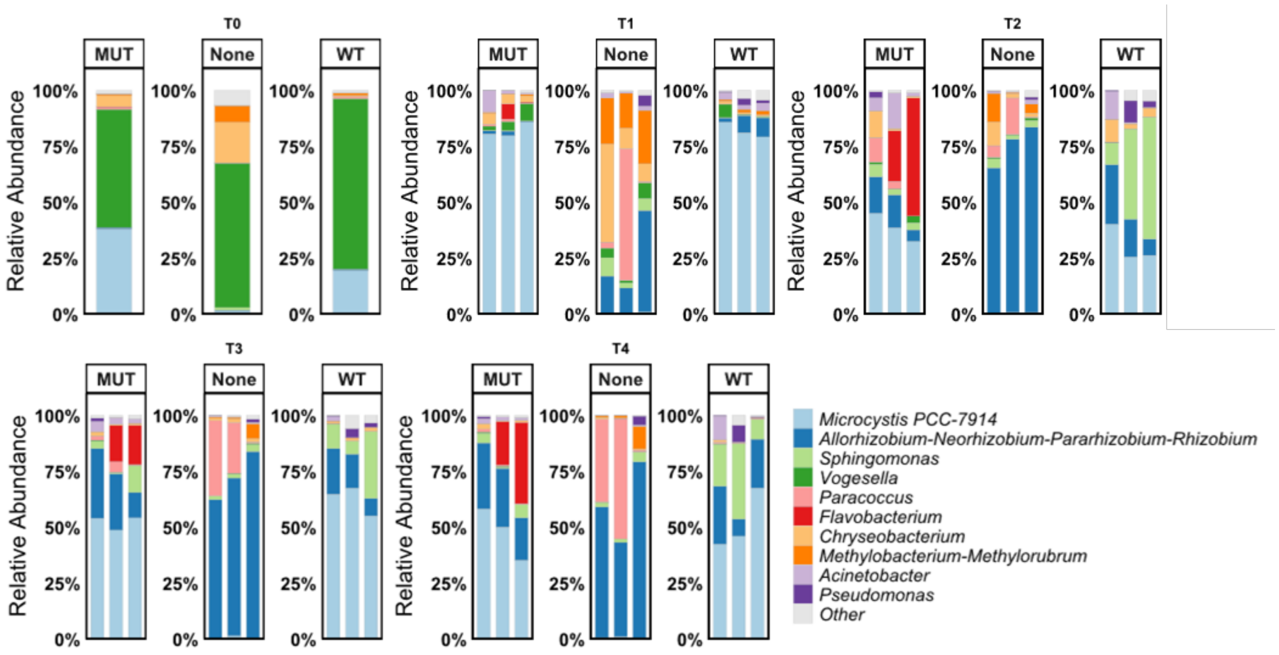

**Figure S4**: Relative abundances of heterotrophic bacterial genera in SynCom experiment. Each bar represents a replicate in the respective group. Sampling time points are indicated above the panels. MUT = co-cultivation with MC- strain, WT = co-cultivation with MC+ strain, None = cultivation of heterotrophic bacterial consortium without cyanobacteria.

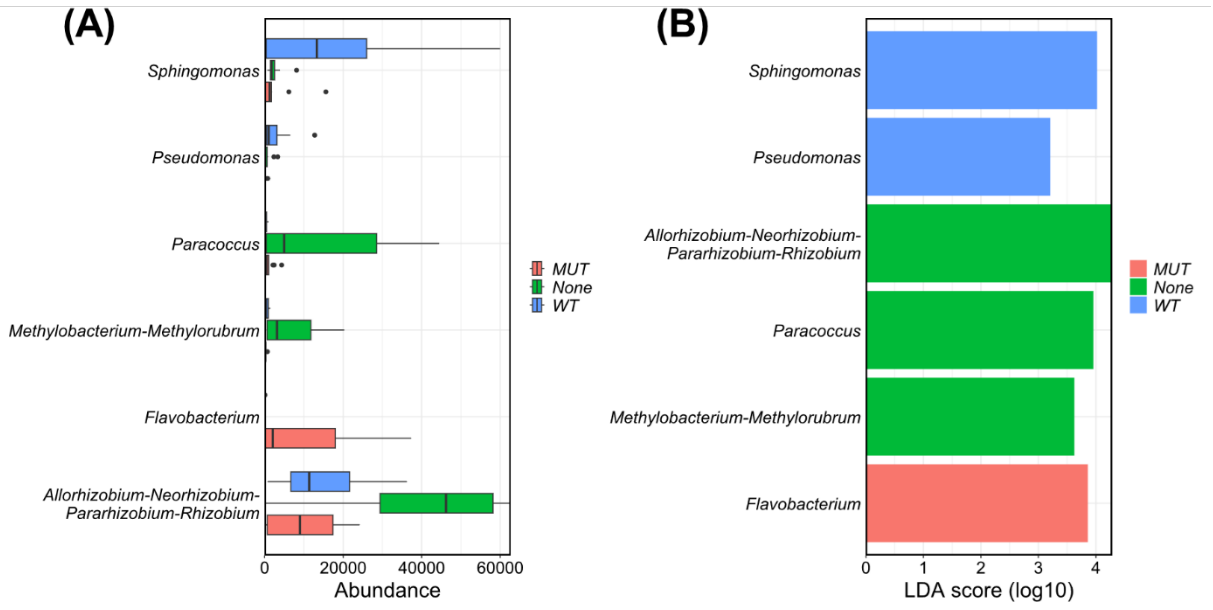

**Figure S5**: LEfSe analysis of SynCom experiment. (A) Boxplot of CSS abundance of taxa scored by LEfSe analysis of synthetic communities. (B) Log10-LEfSe Score of taxa with cutoff = 3.

### *Agrobacterium*

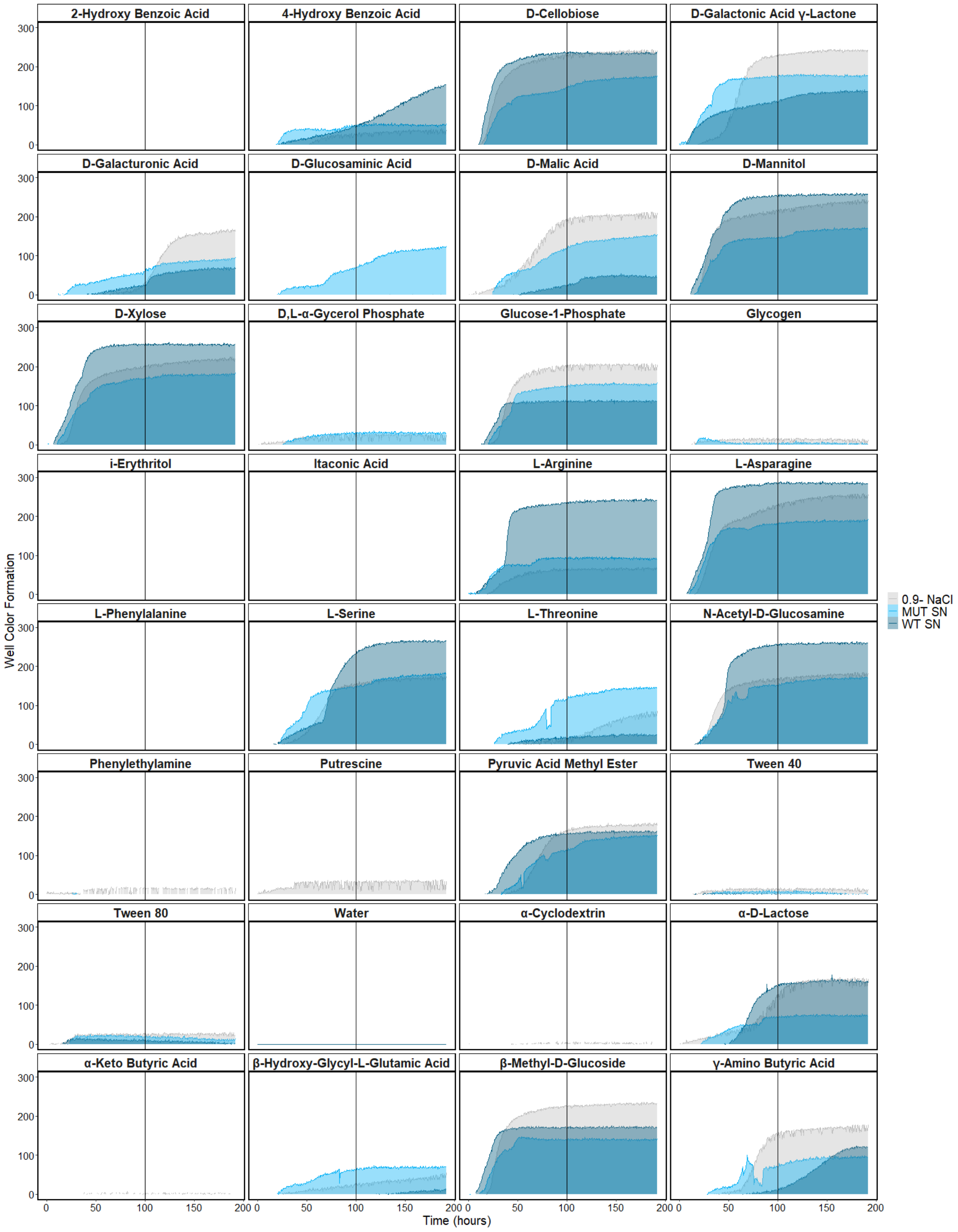

**Figure S6**: EcoPlate™ substrate utilization test of *Agrobacterium* sp. UP1. Well color was measured as an indicator of metabolic activity by Biolog Software every 30 min for 192 hours. *Agrobacterium* was resuspended either in *Microcystis* culture exudates (WT exudates: dark blue, MUT (Δ*mcyB*) exudates: light blue) or 0,9%-NaCl solution (grey). Vertical line indicates the datapoint used for the heatmap (Fig. 4)

### *Sphingomonas*

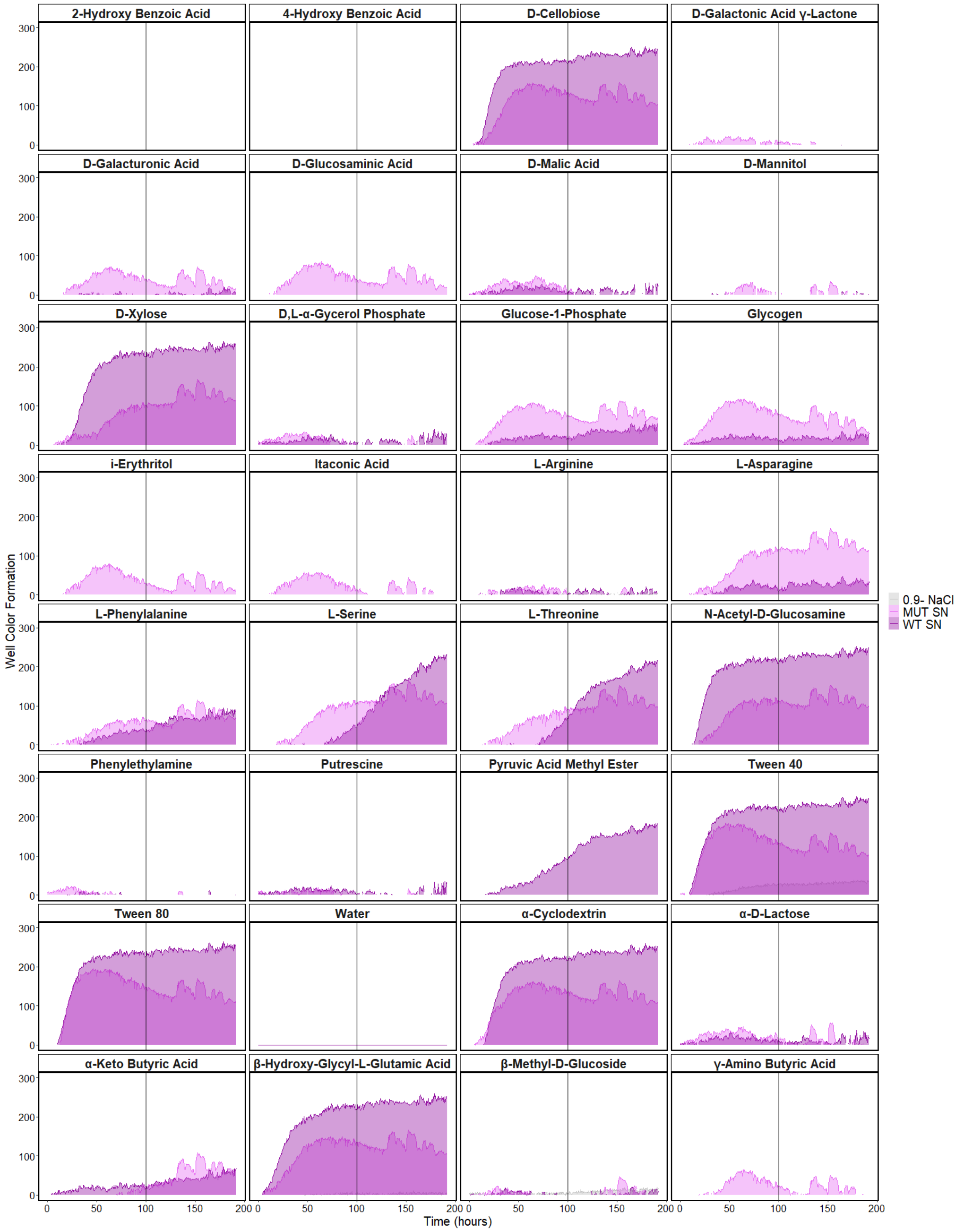

**Figure S7**: EcoPlate$^{TM}$ substrate utilization test *Sphingomonas* sp. UP3. Well color was measured as an indicator of metabolic activity by Biolog Software every 30 min for 192 hours. *Sphingomonas* was resuspended either in Microcystis culture exudates (WT exudates: dark pink, MUT (Δ*mcyB*) exudates: light pink) or 0,9%-NaCl solution (grey). Vertical line indicates the datapoint used for the heatmap (Fig. 4)

### *Flavobacterium*

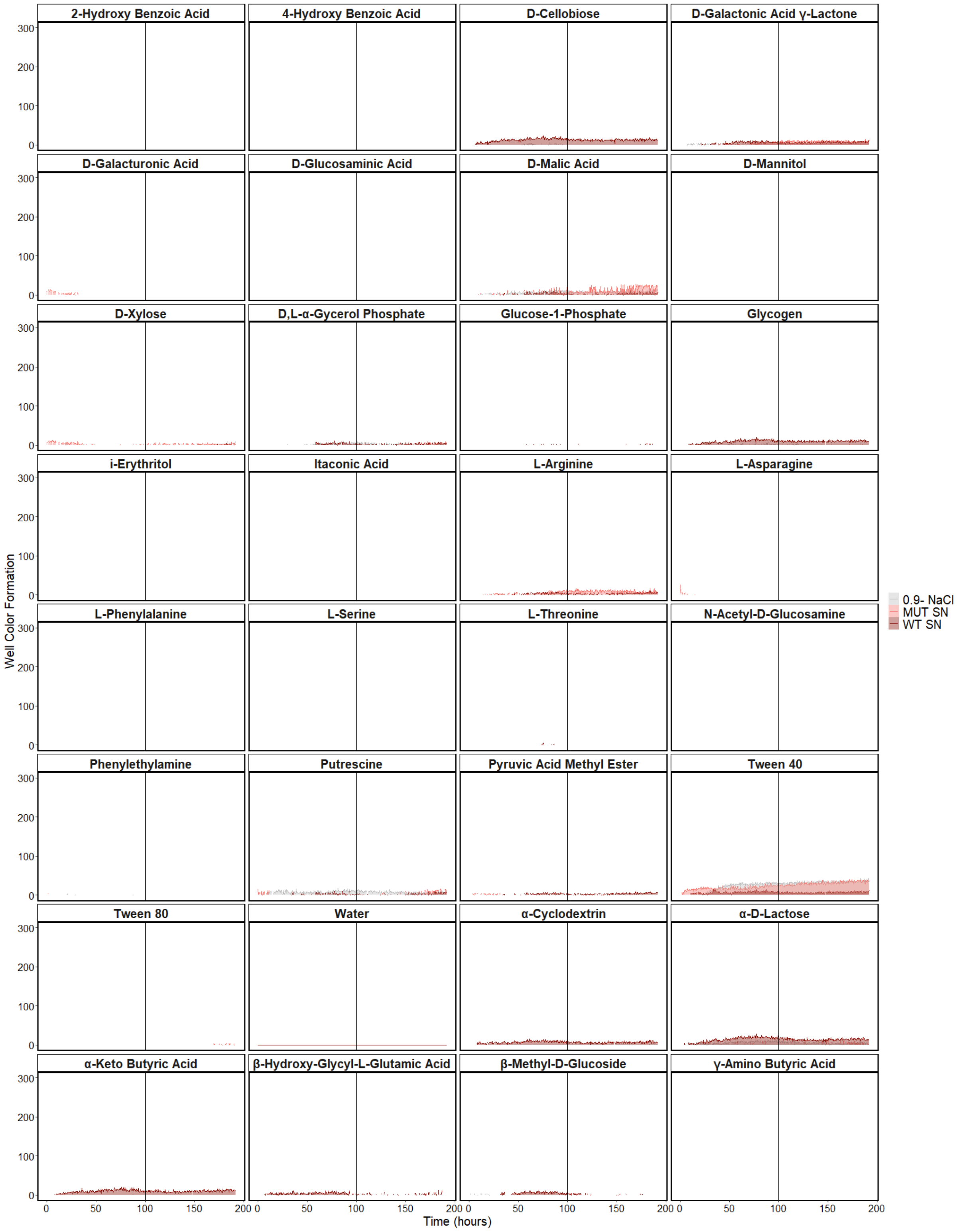

**Figure S8**: EcoPlate™ substrate utilization test of *Flavobacterium* sp. UP2. Well color was measured as an indicator of metabolic activity by Biolog Software every 30 min for 192 hours. *Flavobacterium* was resuspended either in *Microcystis* culture exudates (WT exudates: dark pink, MUT (Δ*mcyB*) exudates: light pink) or 0,9%-NaCl solution (grey). Vertical line indicates the datapoint used for the heatmap (Fig. 4)
